# A non-human primate model of familial Alzheimer’s disease

**DOI:** 10.1101/2020.08.24.264259

**Authors:** Kenya Sato, Hiroki Sasaguri, Wakako Kumita, Takashi Inoue, Yoko Kurotaki, Kenichi Nagata, Naomi Mihira, Kaori Sato, Tetsushi Sakuma, Takashi Yamamoto, Michihira Tagami, Riichiroh Manabe, Kokoro Ozaki, Yasushi Okazaki, Takaomi C. Saido, Erika Sasaki

## Abstract

Alzheimer’s disease (AD) is a major cause of dementia, with the number of patients with this condition anticipated to exceed 50 million worldwide in the near future. Despite extensive research efforts, no effective measures are available to facilitate the prevention or treatment of AD, which is due in part to a lack of animal models able to closely replicate a human-like disease state. Here, we describe the generation of three mutant marmoset individuals in which exon 9 of *PSEN1* gene product has been deleted (*PSEN1*-ΔE9). Such ΔE9 mutations have been reported to cause early on-set familial AD (references^1–5^). We used Transcription Activator-Like Effector Nuclease (TALEN) to destroy the 3’ splice site of exon 9 in the marmoset *PSEN1* gene. To this end, TALEN exhibits high genome-editing efficacy, generates few off-target effects, and produces minimal mosaicism. Indeed, whole genome sequencing and other analyses illustrated an absence of off-target effects and an apparent absence of mosaicism. Fibroblasts obtained from newborn marmosets exhibited uncleaved full-length presenilin 1 protein (PS1) caused by the perturbation of PS1 endoproteolysis as well as an increased ratio of Aβ_42_/Aβ_40_ production, a signature of familial AD pathogenesis. To our knowledge, this is the first non-human primate model of familial AD. We intend to make our marmoset model available to the research community to facilitate the global fight against AD.

## Introduction

Alzheimer’s disease (AD) is the most common neurodegenerative disease in humans, depriving patients of their dignity and having an enormous social impact. The number of individuals with dementia in the world was approximately 50 million in 2019^6^, with AD accounting for 50–70% of these cases^7^. Clinically, AD is characterized by early memory deficits followed by a decline in other cognitive functions^7,8^. The pathological changes associated with AD development precede the clinical manifestation by approximately 20 years^9^ and comprise the following: deposition of amyloid β peptide (Aβ) as extracellular plaques, accumulation of hyper-phosphorylated tau as intracellular neurofibrillary tangles (NFTs), and chronic neuroinflammation followed by neurodegeneration mainly in the cerebral cortex and hippocampus^10,11^. Genetic and pathological observations collectively support the Aβ hypothesis depicting that Aβ plays a central role in the AD pathogenesis^12^. Mouse AD models appear to have reached a highly advanced state of development in recent times by overcoming overexpression artefacts^13–17^. The amyloid precursor protein *(App)* knock-in mice that harbor familial AD mutations with the humanized Aβ sequence^13^ reproduce the Aβ pathology and neuroinflammation without overexpression of APP. These mice exhibited cognitive dysfunctions as analyzed by Intellicage,^18^ which occurred presumably as a consequence of vasoconstriction by pericytes^19^, impairment of grid cells,^16^ and a plaque-induced altered gene network^17^, thereby providing mechanistic insights into the action of Aβ pathology.

These mouse models, however, did not exhibit tau pathology or neurodegeneration even when crossbred with human *MAPT* knock-in mice, in which the entire *Mapt* gene had been humanized^15,20^. The reason for the absence of tauopathy and neurodegeneration in these animals remains elusive, but it may simply be because mice only live for approximately two years whereas pathological changes in the human brain proceed over a course of decades^9^. The discrepancy between mice and humans may also be accounted for by species differences in genetics, neuroanatomy, immunity, and metabolism. In addition, various higher cognitive functions specific to primates, represented by the presence of a highly developed prefrontal cortex^21^, are affected in AD. In this respect, the mouse models are more suitable for preclinical than for clinical studies. We thus came to the conclusion that establishing a non-human primate model for “near”-clinical studies is a clear research priority^22–24^.

Common marmosets (marmosets, *Callithrix jacchus*) are small, non-human primates that belong to the New World primate family. They are being used increasingly in neuroscience because of advantages over other research primates that have become evident^22^ **(Supplementary Table 1**). Marmosets possess genetic backgrounds, physiological functions, brain structures and complex cognitive/social behaviors resembling those of humans; they communicate mainly via visual and auditory measures. In relation to AD research, the amino acid sequence of Aβ in marmosets is identical to that of humans, with wild-type marmosets starting to accumulate Aβ from 7 years of age or even earlier^25,26^. In addition, adolescent marmosets exhibit tau hyperphosphorylation, but not NFT formation, in the brain that increases with aging^26^. The life span of marmosets in captivity can be as long as 10 to 15 years, making these animals a suitable model for research into aging^27^. As their immune systems and metabolic functions resemble those of humans,^27,28^ the pathogenic processes related to AD may thus be similarly affected^29–31^. Because sleep disorder is an early clinical symptom of AD^32^, it is noteworthy that marmosets share similar sleep phases with humans, as evidenced by the presence of rapid eye movement (REM) and non-REM cycles^33^. Among various non-human primate species, the marmoset seems most applicable to genetic manipulation, i.e. generation of designed mutants, for which their high reproductive efficacy is advantageous^22,34^. Furthermore, fecundity characteristics of marmosets, such as their short period to reach sexual maturity, multiple births, and short gestation interval, are suitable for the production of genetically modified disease-based models. We opted to introduce a pathogenic mutation in the marmoset *PSEN1* gene because the majority of familial AD-causing mutations reside in the *PSEN1* gene^35^. Typically, deletion mutations in exon 9^1–5^ or point mutations at the 3’ splice site (acceptor site) of exon 9 in the *PSEN1* gene cause dominantly inherited familial AD. The point mutations instigate exon 9 elimination and S290C modification in the corresponding mRNA at the junction sites of exons 8 and 10 via conversion of alternative splicing^36–40^. We thus set out to generate a marmoset model of AD in which exon 9 of *PSEN1* gene product is deleted using TALEN to produce AD in these animals. TALEN exhibits high genome-editing efficacy, generates few off-target effects, and produces minimal mosaicism^34,41^, thus making this approach highly useful for the generation of a marmoset model of AD.

## Results

### Evaluation of TALEN activity

We used the Platinum TALENs designed to target the 3’ splice site of exon 9 of the marmoset *PSEN1* gene (**Figure 1a**)^42,43^. After introduction of the TALEN mRNAs into the nuclei of marmoset pronuclear stage embryos, we performed surveyor assays and sequencing with genomic DNA extracted from developed embryos. We found that two out of three embryos exhibited deletions at the target sequences including the acceptor site as expected (**Figure 1b**). To confirm the exclusion of exon 9 in the *PSEN1* mRNA, we performed RT-PCR and sequenced the corresponding cDNA sequencing using RNAs extracted from 4-cell-stage or single blastomeres of the TALEN-injected marmoset embryos (**Figure 1c**)^34^. We consequently verified complete exclusion of exon 9 in two out of two 4-cell-stage embryos and three out of five single blastomeres failed to amplify cDNA (**Figure 1d**). These findings indicated that the 3 ‘ splice site-deletion resulted in the exclusion of exon 9 in the mRNA transcribed from the *PSEN1* gene. There was no wild-type sequence in the embryos or single blastomeres in which the 3’ splice site was destroyed, suggesting that the mutations took place in a biallelic manner. We later noted that homozygous deletion of *PSEN1* exon 9 appeared to cause embryonic lethality *in vivo* (see below).

**Figure 1.**
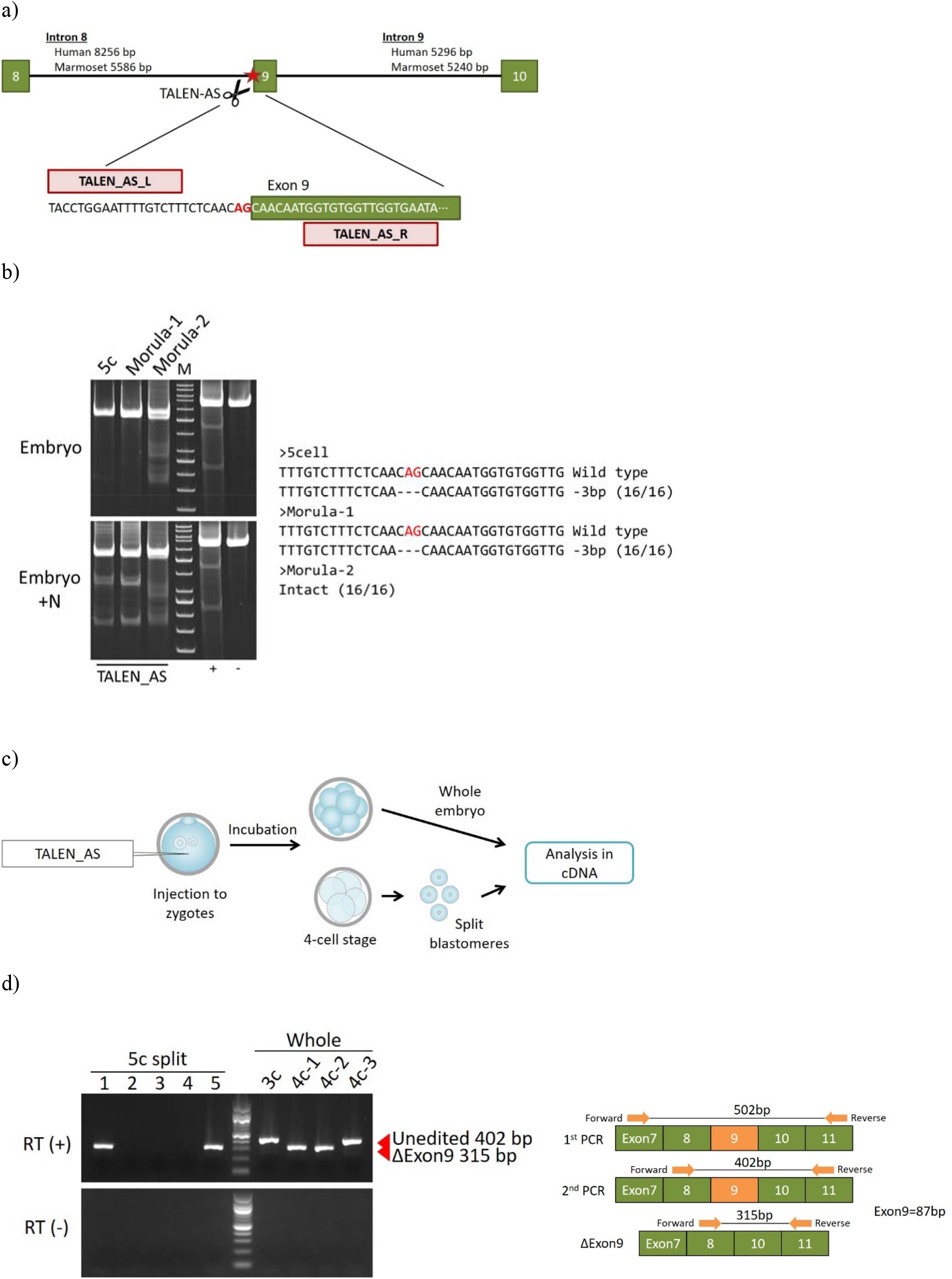
TALEN-mediated genome editing in marmoset embryos. **a)** Left (TALEN_AS_L) and right (TALEN_AS_R) were constructed using the Platinum Gate TALEN Kit (Addgene; cat#10000000439) as previously described^43^. Pink boxes indicate TALEN targeting sites, green boxes indicate exons of the marmoset *PSEN1* gene, and red letters (AG) represent the acceptor site of exon 9. **b**) Surveyor assay of marmoset embryos after TALEN injection. Samples, in which the PCR product were mixed in equal amounts with the PCR product from wild-type *PSEN1* DNA as a template to detect biallelic mutations in the *PSEN1* gene, are designated as +N (three leftmost columns) was performed with or without nuclease (N) treatment. The right two columns denote experimental controls included in the Surveyor mutation detection kit. M; GeneRuler 50bp DNA Ladder (Thermo scientific, SM0371), +N; Equal amount of normal *PSEN1* PCR products added, +; Positive control in Surveyor mutation detection kit, -; Negative control in Surveyor mutation detection kit. **c)** Schematic representation of TALEN mRNA injection and subsequent analysis of *PSEN1* mRNA. After injection of TALEN mRNA into marmoset zygotes, RNA was extracted from four 3- to 4-cell stage whole embryos or single blastomeres from one 5-cell stage embryo after blastomere splitting. The RNA was reverse transcribed, and the cDNA was subjected to PCR (RT-PCR). **d)** RT-PCR of mRNA derived from TALEN-injected embryos. Lanes 1-5 indicate RT-PCR products of each blastomere after splitting; lanes 3c, 4c-1, 4c-2, and 4c-3 indicate RT-PCR of four embryos without splitting. PCR products with exclusion of exon 9 appeared as 315 bp bands while wildtype bands were 402 bp. Note that all the analyzed embryos showed only a single band, suggesting that no embryos carried the mutations in a heterozygous manner. M; 100bp DNA Ladder (New England Biolabs; N3231), RT (+); Reverse transcriptase-treated samples, RT (-); None-treated sample instead of reverse transcriptase.

### Generation of marmosets lacking *PSEN1* exon 9 in the corresponding mRNA (*PSEN1*-ΔE9) and apparent absence of mosaicism

The embryos treated with TALEN were transferred into the uteri of surrogate mother marmosets to generate *PSEN1* exon 9-deleted AD model marmosets. However, the developmental rate of the embryos (35%) was significantly lower than that in our previous study^34^, and no pregnant animals were obtained even after 40 zygotic injections. These results implied that the deletion mutations of *PSEN1*-ΔE9 in the biallelic status are likely to cause the low embryonic developmental rate and embryonic lethality, presumably due to the disruption of Notch signaling^44^. This was somewhat expected because mice deficient in PS1^45,46^ or lacking active site aspartate^47^ exhibited embryonic lethality and because the protein domain corresponding to *PSEN1* exon 9 is a site for endoproteolysis^48^, which is perturbed by destruction of the active site^47^. We thus came to realize that the ΔE9 mutations needed to be introduced in a monoallelic manner.

To avoid biallelism of the *PSEN1* gene mutations, we injected TALEN mRNAs into 810 ova and then performed *in vitro* fertilization on the 578 (71.4%) ova that survived, using wild-type marmoset sperm. After the *in vitro* fertilization, 218 (37.7%) zygotes were obtained, following which 154 (70.6%) embryos that developed to above the 6-cell stage were transferred into the uteri of 77 surrogate mothers. Around the 145th day from the ovulation of the surrogate mother, six marmoset neonates were obtained by normal delivery or by caesarean section (**Figure 2a, Table 1**).

**Figure 2.**
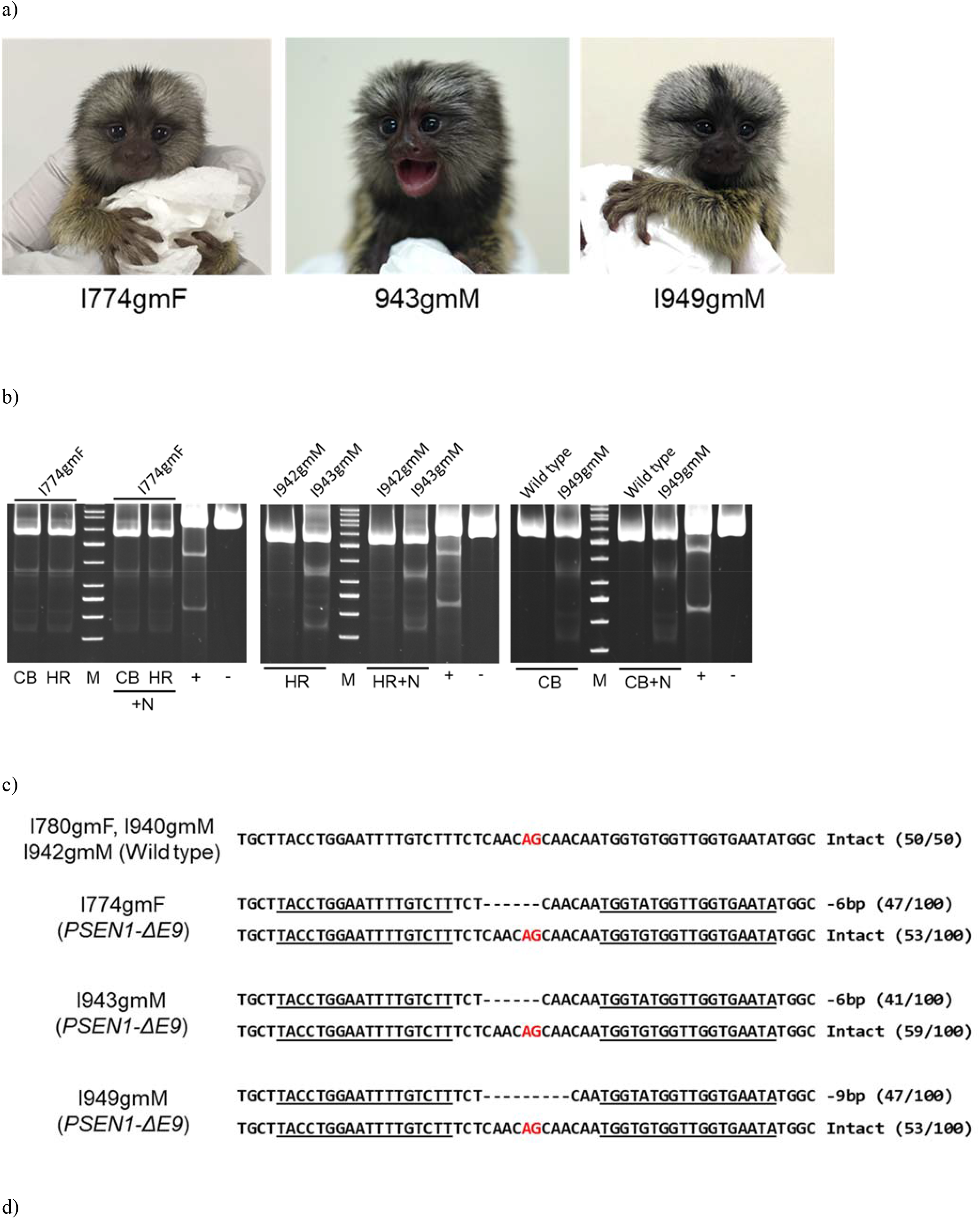
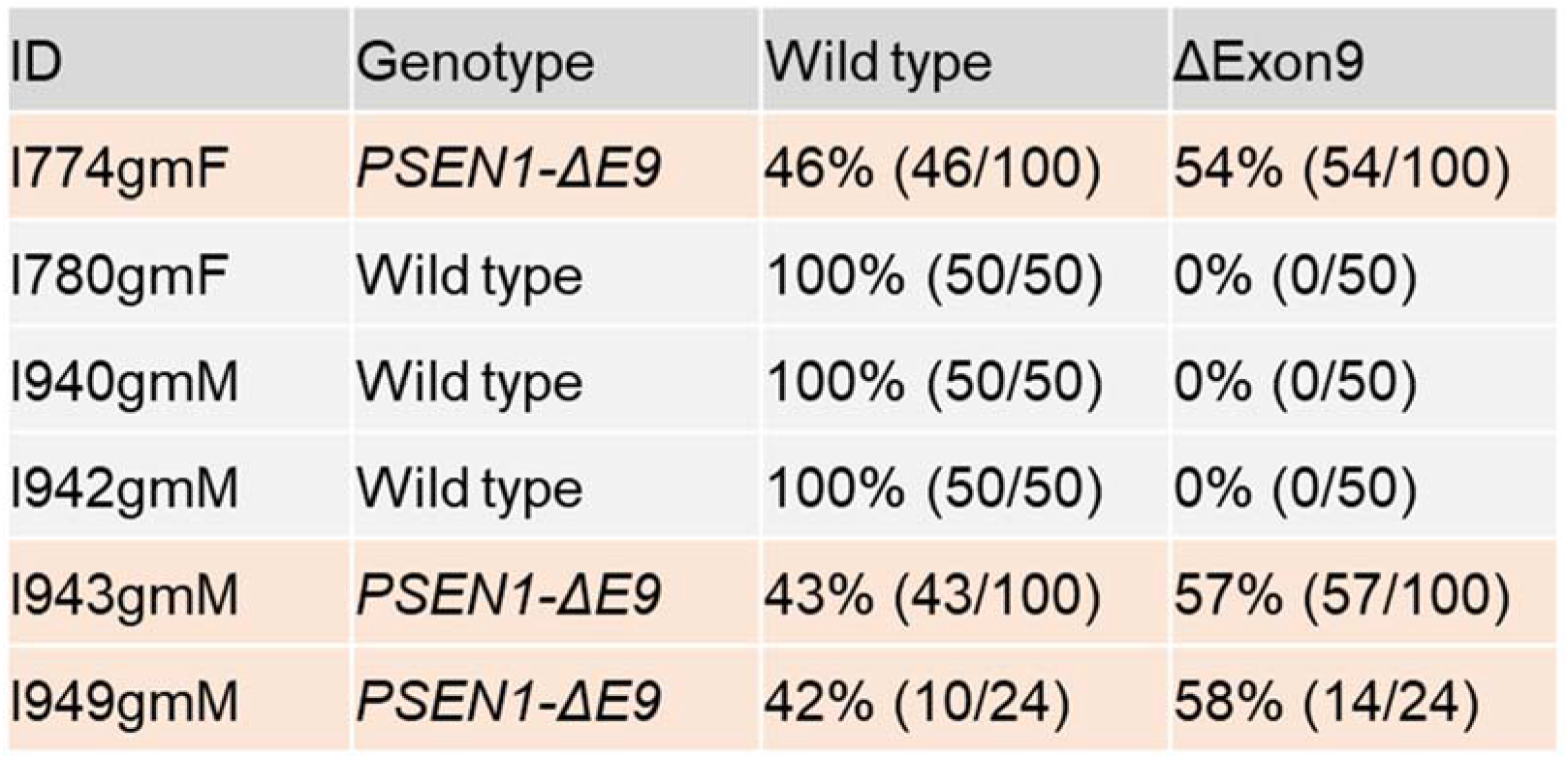
The *PSEN1-ΔE9* marmosets generated by TALEN. **a)** Images of founder neonates. **b)** Surveyor assays of neonatal somatic tissues. Genomic DNAs extracted from umbilical cord blood and hair roots were subjected to surveyor assays. CB; Cord blood genome, HR; Hair root genome, M; GeneRuler 50bp DNA Ladder, +N; Equal amount of normal PSEN1 PCR products added, +; Positive control in Surveyor mutation detection kit, -; Negative control in Surveyor mutation detection kit. **c)** Sequences of the TALEN-targeted site in *PSEN1* gene. The underlined section corresponds to the TALEN recognition sequence: the red letters to the acceptor site, and the dashes to the missing bases. **d)** Results of sequence analyses of neonates. RT-PCR products from hair roots obtained from six neonates were subcloned and sequenced. The first column shows the marmoset identities, the second their genotypes, the third the percentage of wild-type cDNA, and the last column the percentage of mutated cDNA.

**Table 1.** Summary of animal generation.

| TALEN<br>injected<br>ova | Matured<br>ova (%) | Fertilized<br>ova (%) | Transferred<br>embryos<br>(%) | Recipients | Pregnant<br>animals<br>(%) | Aborted<br>case<br>(%) | Delivered<br>animals | Intact | Mutant |
| --- | --- | --- | --- | --- | --- | --- | --- | --- | --- |
| 810 | 578<br>(71.4) | 218<br>(37.7) | 154 (70.6) | 77 | 7 (9.1) | 2 (28.6) | 6 | 3 | 3 |

Surveyor assays of the genomic DNA isolated from hair roots of the newborns suggested that three out of six neonates carried *PSEN1* gene mutations (**Figure 2b**). Sequence analyses of the subcloned PCR products from the genomic DNA indicated three different patterns of *PSEN1* gene mutations in these animals. Two animals carried 6-bp deletions, while another newborn harbored 9-bp deletions at the 3’ splicing site, indicative of’ deletion of the region in mRNA corresponding to the genomic *PSEN1* exon 9 (**Figure 2C**). The numbers of wild-type and truncated genes in the subclones collected from the mutants’ hair roots were 53 and 47 out of 100 in I774gmF, 51 and 49 in I943gmM, and 59 and 41 in I949gmM, respectively (**Figure 2d**): approximately 50% of the clones carried identical mutations in each case. Consistently, RT-PCR followed by cDNA sequencing of total RNA from hair root cells verified the excision of exon 9 in the mRNAs extracted from hair root samples of the mutants. These observations suggest two possibilities: [1] the presence of heterozygous mutations essentially in all cells; and [2] the presence of wild-type and homozygous mutant cells in a 1:1 ratio, i.e. mosaicism. The second possibility can be ruled out because homozygous *PSEN1*-ΔE9 mutations result in embryonic lethality, presumably via a Notch phenotype^5,46 49^ that would preclude half of the neural stem cells from differentiating to neurons during development. It is noteworthy that γ-secretase cleaves the truncated Notch in a cell-autonomous manner^50^. These results and other speculations^34^ indicate a probable absence of mosaicism in the mutant marmosets. The sequence analyses of the cDNAs also showed that there were no insertions or deletions in the exons 8 or 10 in the *PSEN1* gene (**Figure 2d, Supplementary Figure 1**). We thus concluded that we had successfully acquired *PSEN1*-ΔE9 marmoset neonates.

### Off-target analyses of the mutant *PSEN1*-ΔE9 marmosets

A number of genome editing-based studies suffer from off-target effects^41^. We checked for off-target mutations in the mutant *PSEN1*-ΔE9 marmosets in two ways. The online Paired Target Finder Tool was used to search for candidate off-target sites^34^ to enable calculation of an average score for each potential off-target site obtained from the search. In this way, the top 10 candidate sites were identified (**Supplementary Table 2**). Genomic DNA extracted from hair roots of *PSEN1*-ΔE9 marmosets was subjected to PCR, and the sequence then determined by direct sequencing. None of the *PSEN1*-ΔE9 marmosets exhibited any mutations in the off-target candidate sites. Second, we performed whole genome sequencing (WGS) of the genomic DNA obtained from the first *PSEN1*-ΔE9 marmoset (I774gmF) and from her parents at an average coverage over the genome of 42.1, 57.4, and 62.2x for the neonate, father, and mother, respectively (**Supplementary Table 3**). In order to process the WGS data sets, we performed Illumina Dynamic Read Analysis for GENomics (DRAGEN) pipeline (version 3.5.7)^51^ for mapping and variant calling process at a default parameter with the “CJA1912RKC” marmoset genome reference sequence (GCA_013373975.1). We used this highly permissive parameter in variant calling and did not filter out variants in order to keep as many variants in off-target candidate sites as possible. By comparing all the variants from the *PSEN1*-ΔE9 marmoset (I774gmF) with those from her parents, we found 306,492 variants only in the *PSEN1*-ΔE9 marmoset (I774gmF) (**Supplementary Figure 2**). None of these variants were present within the 10 off-target candidate sites, further confirming the absence of off-target mutations. We thus concluded that application of the TALEN technology to generate the *PSEN1*-ΔE9 marmosets has fully succeeded.

### Endoproteolysis of mutant marmoset-derived PS1 and secretion of Aβ_42_ versus Aβ_40_

PS1 serves as a major catalytic subunit for the γ-secretase complex that generates Aβ from the C-terminal fragment of APP after limited proteolysis by β-secretase^45,47,48,52^. Most, if not all, of the pathogenic mutations found in the *PSEN1* gene cause familial AD by elevating the ratio of Aβ_42_/Aβ_40_ production^53–55^. The ratio matters because, as demonstrated by Kim *et al.,*^56^ Aβ_42_ is causal whereas Aβ_40_ is protective in the pathological processes. To biochemically analyze the mutant marmosets in the least invasive manner, we established primary fibroblasts from tissue excised from edge of the ear lobe in wild-type and mutant animals. We first examined the effect of exon 9 deletion on the endoproteolytic status of PS1 protein because such a deletion impacts on the global PS1 endopeptidase activity^5^. We performed Western blot analysis of fibroblasts derived from wild-type and *PSEN1*-ΔE9 marmosets using a set of antibodies that recognize the N- and C-termini of PS1. The exon 9 deficiency in the two independent mutants, i.e. I774gmF and I1943gmM, gave rise to a full-length PS1 that was absent in the wild-type fibroblasts (**Fig. 3a**). Consistently, the quantities of N-terminal and C-terminal fragments in the mutants were reduced to approximately half the levels of those found in wild-type cells. These observations indicate that the monoallelic exon 9 deletion caused perturbation of the PS1 endoproteolysis in a heterozygous manner and that the nature of endoproteolysis was autolytic. The deletion also induced an amino acid sequence conversion, S290C, as shown in **Supplementary Figure 1**^40^.

**Figure 3.**
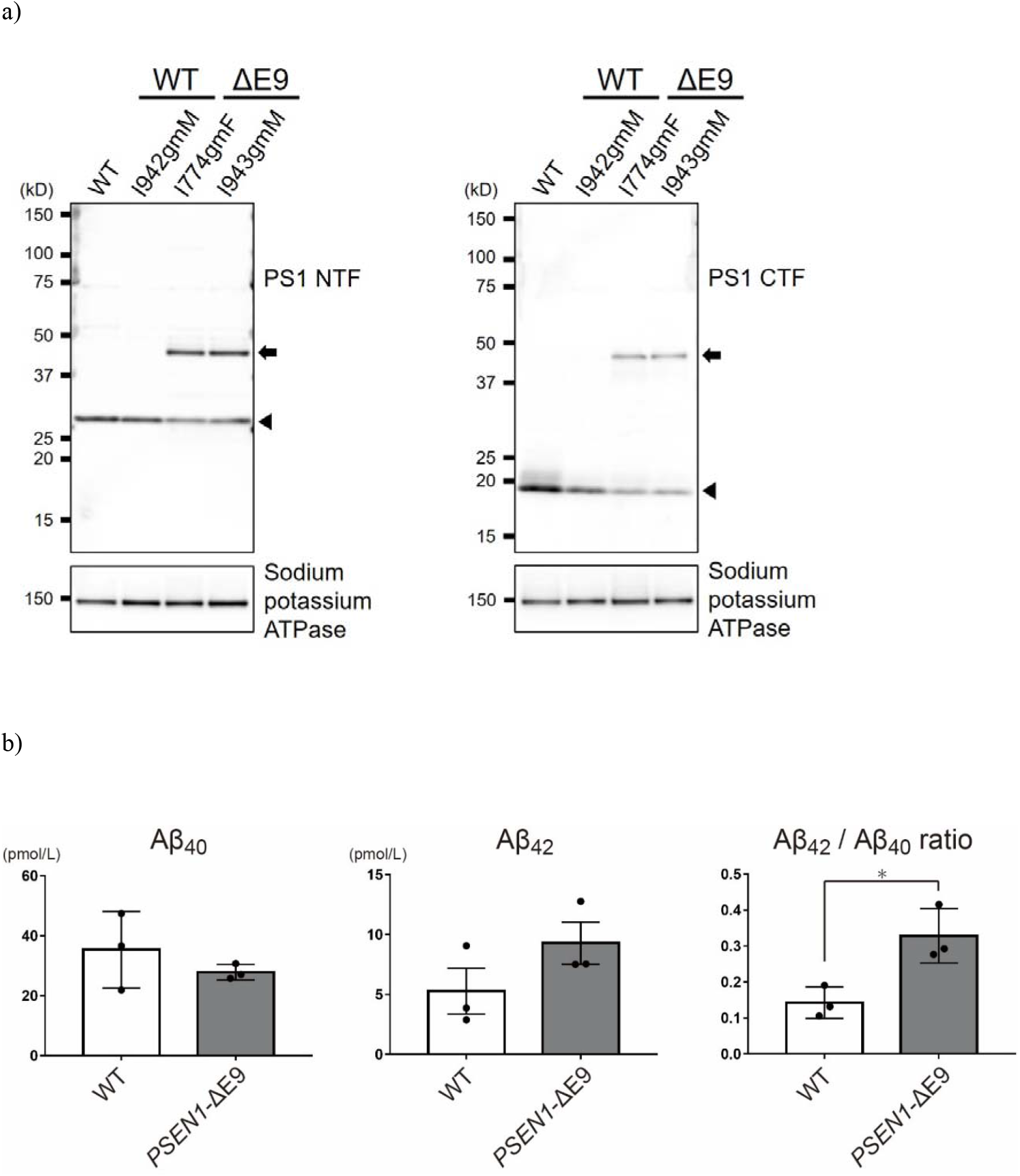
Biochemical analysis of *PSEN1-ΔE9* marmoset fibroblasts. **a)** Western blot analysis of the membrane fractions obtained from wild-type (WT) and mutant (ΔE9) marmoset fibroblasts using antibodies to N-terminal fragments (NTF) (left panel) and to C-terminal fragments (CTF) (right panel) of PS1. Arrowheads indicate NTF and CTF produced by endoproteolysis of PS1; arrows indicate uncleaved, full-length PS1 protein. Sodium-potassium ATPase was used as a loading control of the membrane fraction. **b)** Aβ_40_ and Aβ_42_ in the culture media from wild-type and mutant marmoset fibroblasts were quantified by sandwich ELISA. Aβ43 levels were below the detection limit. Data represent mean ± sem (n=3). WT; wildtype. \**P* ≤ 0.05 (Student’s two-tailed *t*-test).

We also quantified Aβ_42_ and Aβ_40_ secreted from the wild-type and mutant fibroblasts by enzyme-linked immunosorbent assay (ELISA) for the reasons stated above. The quantity of Aβ_42_ was significantly increased in the mutant-derived media while that of Aβ_40_ showed a tendency towards reduction. Consequently, the ratio of Aβ_42_/Aβ_40_ production was statistically significantly increased more than 2-fold i (**Fig. 3b**), a hallmark of familial AD pathogenesis. Similar results have been reported using cultured human fibroblasts and lymphocytes obtained from *PSEN1* mutation carriers^53–55^. Furthermore, Aβ_43_^57^ was undetectable in the present experiments. Taken together, these observations indicate that the *PSEN1*-ΔE9 mutations introduced into the marmosets are indeed pathogenic.

## Discussion

Steiner *et al.* previously demonstrated that a *PSEN1*-ΔE9 mutation exerted its pathogenic effect via an amino acid conversion, S290C, rather than via perturbed PS1 endoproteolysis^40^. This effect of the amino acid conversion on the Aβ_42_/Aβ_40_ ratio is also likely to apply to our observation concerning fibroblasts obtained from mutant marmosets since they also carry the S290C conversion (**Supplementary Figure 1**). There is, however, a distinct difference between the studies by Steiner *et al.* and by us. Steiner *et al.* used a cDNA overexpression paradigm in cultured cells and in *Caenorhabditis elegans.* This approach does not necessarily warrant the stoichiometric formation of the γ-secretase complex,^52^ and is comparable to homozygous deletion of exon 9 *in vivo*, which resulted in embryonic lethality (see **Generation of marmosets lacking *PSEN1* exon 9 in the corresponding mRNA (*PSEN1*-ΔE9) and apparent absence of mosaicism**). In addition, the results shown by Steiner *et al.* do not exclude the possible need for the simultaneous presence of exon 9 deletion and S290C mutation together to evoke the pathogenic activity in a coordinated manner. Moreover, the S290C mutation by itself is unlikely to account for the symptom of spastic paraparesis often observed in *PSEN1*-ΔE9 patients^1^. The embryonic lethality caused by the biallelic presence of *PSEN1*-ΔE9 mutations also calls into question the pathogenic role of the S290C mutation by itself because most pathogenic point mutations in the *Psen1* gene generally result in non-lethal phenotypes in a homozygous status, at least in mice, with the exception of a few cases^57,58^. We thus cannot rule out the possible role of ΔE9 mutations alone in the pathogenicity *in vivo*. These issues would be better addressed if we could create various mutant knock-in mouse phenotypes; however, deletion of *Psen1* exon 9 in mice generated a termination codon within the *Psen1*-coding sequences (Kaori Sato and Nagata, unpublished observations), rendering such attempts difficult. While these queries remain as open questions, they are beyond the scope of our present study whose singular aim was to establish a non-human primate model of familial AD.

Currently, the CRISPR/Cas9 technology including Base Editor (BE) appears to be the prevalent method for genome editing in general. Indeed, Wang *et al.* succeeded in generating a Hutchinson–Gilford progeria syndrome cynomolgus monkey model using BE^59^. In the present study, we used a different technique – TALEN – because it is highly efficient, and produces less mosaicism^34^. Furthermore, the application of BE to genome editing of the marmoset *PSEN1* gene has proven to be more challenging than expected (Kumita *et al.,* unpublished observations). The genome editing efficacy by TALEN in the present study was consistently as high as 50%. This study showed that targeting the 3 ‘ splice site of the *PSEN1* gene induced whole exon 9 skipping by TALEN, with cDNA sequence analyses showing that neither insertion nor deletion occurred at exon 8 or exon 10 in the *PSEN1* gene. The strategy used in this study – to induce mutation(s) at the splice site of the target gene – is a new method for the precision deletion of a whole target exon at the target gene, without any mutation in the neighboring exon. Therefore, destruction of the splice site in the *PSEN1* gene to induce exon 9 skipping by this specific genome-editing tool appears to be the most realistic approach to generate a non-human primate model of familial AD. The oldest mutant marmoset among the three mutants, I774gmF, was born in April of 2019 and therefore is still sexually immature, making it difficult to provide direct evidence of the germline transmission of the pathogenic mutations. However, we presume that the apparent lack of mosaicism (see **Results**) implies the presence of the mutations in the reproductive cells because a similar approach to generate IL2RG knock-out marmosets using TALEN resulted in 100% germline transmission^34^.

To our knowledge, this is the first non-human primate model of familial AD ever produced. Given that the age-of-onset of AD in *PSEN1*-ΔE9 humans commonly occurs in the fourth decade^1 2 39^, with Aβ deposition commencing approximately two decades prior to the disease onset in these patients^9^, and that wild-type marmosets start accumulating Aβ in the brain by around the age of 7 years or even earlier^25^, we anticipate the mutant marmosets to start exhibiting Aβ pathology from 2-3 years of age. It should be possible to verify the relevance of the mechanistic findings obtained using model mice^16,17,19^ if the mutant marmosets prove to reflect human AD in a reliable manner. Because marmosets live much longer than mice, they may develop tau pathology and neurodegeneration, which model mice fail to recapitulate. If so, marmosets are likely to become a convenient model to establish cause-and-effect relationships and to elucidate the mechanisms by which Aβ amyloidosis evokes tau pathology and neurodegeneration. Use of the marmoset model could be applicable in the first instance to basic studies necessary for disease verification in humans, including non-invasive imaging analyses by MRI to determine morphometric changes and by positron emission tomography (PET) to search for Aβ and tau pathology. Body fluids such as blood and cerebrospinal fluid may also be collected from marmosets for biomarker identification and development. In a more clinical sense, the marmosets can be subjected to highly sensitive cognitive analyses such as touch-sensitive technology^60^ or for the analysis of sleep disturbances^33^. Functional MRI could also be used to examine the default mode network, changes to which represent an early sign of preclinical AD^61,62^. To this end, model mice are not suitable for such studies, or for understanding the effect of Aβ pathology on prefrontal cortex functionality,^21^ which can only be understood by analyzing the mutant marmosets. For pharmaceutical studies, the mouse model can be used in the first instance for initial screening, with the marmoset model then used for subsequent screening prior to clinical trials. The marmosets could also enable examination of the efficacy of anti-tau pathology and anti-neurodegeneration medication candidates if these pathological features are shown to be present. We are making maximum efforts to expand our colonies so that this marmoset model can be shared with the global AD research community in the near future.

## Methods

### Animals

All animal experiments were carried out in accordance with guidelines for animal experimentation from the RIKEN Center for Brain Science and the Central Institute for Experimental Animals (CIEA: 18031A, 19033A). The CIEA standard guidelines are in accordance with the guidelines for the proper conduct of animal experiments as determined by the Science Council of Japan. The marmosets used in this study were purchased from CLEA Japan, Inc., Tokyo, Japan. The animals ranged from 2-4.5 years of age with body weights between 300 and 450 g. All animals were housed in cages in accordance with the following two international standards for laboratory animal care; National Research Council of the National Academies; Guide for the Care and Use of Laboratory animals (USA), EU commission recommendation of 18 June 2007 on guidelines for the accommodation and care of animals used for experimental and other scientific purposes (EU). The cage consisted of one main body (W900mm x D750mm x H2050mm) and two ceiling cages (W300mm x D425mm x H375mm), and marmosets were bred in family units. The light-dark cycle was 12 hours, and CMS-1M (CLEA Japan), which is a complete diet, was ready for free feeding.

### *In vitro* transcription of TALEN mRNA

We employed a two-step Golden Gate assembly method using the Platinum Gate TALEN Kit (Addgene; cat#1000000043) to construct Platinum TALEN plasmids containing the homodimer-type FokI nuclease domain. The assembled repeat arrays were subsequently inserted into the final destination vector, ptCMV-163/63-VR. TALEN mRNA was synthesized using the mMESSAGE mMACHINE T7 Ultra Transcription Kit (Thermo Fisher Scientific, AM1345). Transcribed mRNA was purified using the MEGAclear Transcription Clean-Up Kit (Thermo Fisher Scientific, AM1908), and nuclease-free water (Thermo Fisher Scientific, AM9937) was used for the elution step and subsequent dilution. For microinjection, an aqueous solution was used in which left TALEN and right TALEN mRNAs were mixed in equal amounts so that the final concentration was 8 ng/μl.

### Procedures for oocyte collection and in vitro fertilization

Oocyte collection and *in vitro* fertilization (IVF) was performed as previously described^22,63^. Briefly, 62 female marmosets whose plasma progesterone levels were under 10 ng/ml were used for the follicular stimulation. To collect oocytes, the selected marmosets were intramuscularly treated with purified human menopausal gonadotrophin and follicle-stimulating hormone (hFSH, 25IU; FOLYRMON-P injection, Fuji Pharma) a total of five times every other day. The day after the last hFSH administration, human chorionic gonadotropin (hCG, 75IU; Gonatropin, ASKA Pharmaceutical) was subsequently administered by intramuscular injection. The hormone-treated female marmosets were pre-anesthetized with 0.04 mg/kg medetomidine (Domitor; Nippon Zenyaku Kogyo), 0.40 mg/kg midazolam (Dormicam; Astellas Pharma) and 0.40 mg/kg butorphanol (Vetorphale; Meiji Seika Pharma). The ovum pickup was performed at 17-20 hours after the hCG injection. During the procedure, the marmosets were anesthetized by isoflurane (Isoflurane Inhalation Solution; Pfizer) inhalation. The collected ova were incubated in Porcine Oocyte Medium (Research Institute for the Functional Peptides, IFP1010P) for 28-32 hours for *in vitro* maturation^64^. For IVF, semen was collected from healthy male marmosets and diluted to 3.6 × 10^6^ sperm/ml, after washing with TYH medium (LSI Medience, DR01031). Insemination was performed via co-incubation of ova with 3.6 × 10^4^ sperm per oocyte for 10-16 hours at 37°C with 5% CO_2_, 5% O_2_, and 90% N_2_.

### Blastomere splitting

TALEN-mRNA-injected embryos that had reached the 4-cell stage and above were directed to blastomere splitting as previously described^34^. Briefly, the embryo specimens were treated with acidified Tyrode’s solution (Origio, 10605000) to lyse the zona pellucida. Naked embryos were transferred into Embryo Biopsy Medium (Origio, 10620010). After several minutes, embryos that exhibited weak adhesion between blastomeres were split into single blastomeres using glass capillaries and washed immediately in droplets of phosphate-buffered saline (PBS) (-) drops. The blastomere was then transferred to a test tube with a small amount of PBS and used for subsequent analyses.

### Embryo transfer

Embryos grown to a 6-cell-stage or higher were transferred to the uteri of surrogate mothers as previously described^22,63^. The method for obtaining newborns was based on natural delivery, but if delivery was judged to be difficult, a caesarean section was performed on day 145 after embryo transfer^34^.

### PCR for genotyping

Genomic DNA was extracted from marmoset cord blood and hair roots using a QIAamp DNA Micro Kit (Qiagen, 56304) according to the manufacturer’s instructions. Embryos and blastomeres were used as templates without any genome extraction processing. PCR was performed using specific primers followed by subsequent sequencing or surveyor assay (**Supplementary Table 4**). Each PCR mixture contained 1X PCR buffer, 0.2 mM of dNTPs, 0.5 mM of each primer, 20 ng of template DNA and 2.5 U of KOD-Plus-Neo (Toyobo, KOD-401) in a total volume of 20 μl. PCR was performed using the following PCR cycling conditions: 1st PCR (for embryos and blastomeres), one cycle of 94°C for 2 min, 45 cycles of 98°C for 10 sec, 60°C for 10 sec and 68°C for 40 sec, one cycle of 68°C for 7 min. 2nd PCR (for all samples), one cycle of 94°C for 2 min, 35 cycles of 98°C for 10 sec, 60°C for 10 sec and 68°C for 30 sec, one cycle of 68°C for 7 min.

### Sequencing analyses

To confirm the sequences of the TALEN target sites, PCR products were subcloned into the Zero Blunt PCR Cloning Kit (Thermo fisher scientific, 44-0302) according to manufacturer’s instructions. The plasmids obtained were transformed with ECOS Competent E. coli DH5α (Nippon gene, 316-06233) and cultured on LB agar containing kanamycin. After amplifying the plasmid using the emerged single colony, it was reacted using M13 primer and a BigDye terminator v3.1 cycle sequence kit (Thermo Fisher Scientific, 4337455), and gene sequences were analyzed by a 3130 Genetic Analyzer (Thermo Fisher Scientific).

### Surveyor assay

The SURVEYOR mutation detection kit (IDT, 706025), a mutation analysis method that identifies mismatches in double-stranded DNA, was used to detect genetic modifications at the TALEN target site. Briefly, 8 μl of the PCR product described above in **PCR for genotyping** was used for the SURVYOR assay. For the analysis of biallelic mutations in the *PSEN1* gene, a sample in which 4 μl of the PCR product was mixed with an equal amount of the PCR product prepared using wild-type *PSEN1* as a template was also prepared. The sample was digested with nuclease and electrophoresed on Novex TBE Gels (10%, 12 well; Thermo Fisher Scientific, EC62752BOX) and then subjected to nucleic acid staining with GelRed (Fujifilm Wako Chemicals, 518-24031).

### RT-PCR

Total RNA was extracted from marmoset embryos, blastomeres, and hair roots using Nucleospin RNA plus XS (Takara, U0990B) according to the manufacturer’s instructions. cDNA was synthesized using ReverTra Ace - *α* - (Toyobo, FSK-101F). In the reaction, the sample was divided into two: one received reverse transcriptase, and the other received the same amount of water as a control to evaluate genomic DNA contamination. PCR was performed using the primers designed on exons (**Supplementary Table 4**). Each PCR mixture contained 1X PCR buffer, 0.2 mM of dNTPs, 0.5 mM of each primer, 20 ng of template DNA and 2.5 U of KOD-Plus-Neo (Toyobo, KOD-401) in a total volume of 20 μl. PCR was performed using the following PCR cycling conditions: 1st PCR (for embryos and blastomeres): one cycle of 94°C for 2 min, 45 cycles of 98°C for 10 sec, 60°C for 10 sec and 68°C for 40 sec, and one cycle of 68°C for 7 min. 2nd PCR (for all samples): one cycle of 94°C for 2 min, 35 cycles of 98°C for 10 sec, 60°C for 10 sec and 68°C for 30 sec, one cycle of 68°C for 7 min.

### Whole-genome sequencing (WGS)

To prepare the genomic DNA for WGS, a small section of ear lobe tissue (approximately 3 mm^3^) was excised from marmosets in a lowly invasive body tissue collection procedure. Genomic DNA extraction from the ear tissue was performed using the QIAamp Micro DNA kit (Qiagen, 56304) according to the manufacturer’s instructions. Briefly, the tissue was incubated in lysis buffer and proteinase K overnight at 56°C, then genomic DNA was collected by column purification with low-TE (Tris-HCl 10 mM, EDTA 0.1 mM, pH 8.0) elution. Whole genome sequencing was performed using Illumina NovaSeq. FASTQ files for all WGS runs are deposited under accession numbers DRR239155, DRR239156, and DRR239157 in the DNA Data Bank of Japan (DDBJ) Sequence Read Archive (DRA). For processing the WGS data sets, Illumina DRAGEN pipeline (version 3.5.7) was performed for mapping and variant calling process at a default parameter with the “CJA1912RKC” marmoset genome reference sequence (GCA_013373975.1) (**Supplementary Table 3**). We used this highly permissive parameter in variant calling and did not filter out variants in an effort to keep as many variants in off-target regions as possible. By comparing all the variants from the mutant *PSEN1* (I774gmF) marmoset with those from her parents, 306,492 variants were found in the mutant *PSEN1* marmoset without any filtering.

### Off-target analysis with the Paired Target Finder Tool

Paired Target Finder (https://tale-nt.cac.cornell.edu/node/add/talef-off-paired) was used to search for potential off-target sites^65^. As the reference, CJA1912RKC (GenBank assembly accession: GCA_013373975.1), which is the whole genome sequence of the marmoset, was set in the search database, and the repeat variable diresidue (RVD) sequence of TALEN was provided as follows: RVD Sequence 1, NI HD HD NG NN NN NI NI NG NG NG NG NN NG HD NG NG; RVD Sequence 2, NI NG NG HD NI HD HD NI NI HD HD NI NG NI HD HD NI. The minimum and maximum lengths of the spacers were set to 10 and 30, respectively. Other parameters were set as recommended. To identify genome regions with predicted off-target sequences from the genome reference sequence in the WGS data, a “GGGenome” ultrafast sequence search site was performed and off-target bed files were produced. (URL https://gggenome.dbcls.jp/en/calJacRKC1912_chrM_d3g2003/) Four hundred and twenty-eight paired oriented off-target sequence sets were found from 10 TALEN off-target sequence sets. After filtering (spacer size < 100bp) with the 428 off-target sequencing sets, 10 paired oriented off-target sequence sets were finally predicted as off-target effect candidates. Genomic DNA extracted from marmoset hair roots using the QIAamp DNA Micro Kit (Qiagen, 56304) was subjected to PCR with specific primers followed by direct sequencing (**Supplementary Table 5**). Each PCR mixture contained 1X KOD One PCR Master Mix (Toyobo, KMM-101), 0.5 mM of each primer and 10 ng of template DNA in a total volume of 20 μl. PCR was performed using the following PCR cycling conditions: 30 cycles of 98°C for 10 sec, 60°C for 5 sec and 68°C for 1 sec. The amplicon was purified using NucleoSpin Gel and PCR Clean-up (Takara Bio, 740609), reacted with one of the PCR primers and the BigDye terminator v3.1 cycle sequence kit (Thermo Fisher Scientific, 4337455), and the gene sequence was identified using a 3130 Genetic Analyzer (Thermo Fisher Scientific).

### Western blot analysis

Primary fibroblast lines were established from ear skin tissue extracted from the wild-type and *PSEN1*-ΔE9 marmosets, i.e. I774gmF and I1943gmM, respectively. Western blotting was performed as previously described^66^. Briefly, cells were lysed and subjected to subcellular fractionation using a ProteoExtract Subcellular Proteome Extraction Kit (Merck, #539790) according to the manufacturer’s instructions. We loaded the membrane fractions onto polyacrylamide gels, electrophoresed them, and blotted them onto PVDF membranes (Merck Millipore), which were then treated with ECL Prime blocking agent (GE Healthcare) and incubated with primary antibodies raised against the PS1 N-terminal (G1Nr5) and C-terminal (G1L3) domains^67 68^. We used an antibody to sodium-potassium ATPase (EP1845Y, Abcam, ab76020) as a loading control of the membrane fraction.

### Enzyme-linked immunosorbent assay (ELISA)

Aβ_40_ and Aβ_42_ were quantified using ELISA as previously described^58^. Briefly, culture media, collected from the primary skin fibroblasts 72 hours after incubation, were mixed with 11 times the volume of 6M guanidine-HCl. We then quantified Aβ_40_ and Aβ_42_ using an Aβ ELISA kit (Wako, 294-62501 and 294-62601) according to the manufacturer’s instructions. Quantification of Aβ43 was performed as previously described^57^.

### Statistical analyses

All data are shown as the mean ± s.e.m. For comparison between two groups, data were analyzed by Student’s *t*-test. All the data were collected and processed in a randomized and blinded manner.

## Author Contributions

K.S., H.S., K.N., T.C.S., and E.S. designed this study. K.S., H.S., W.K., N.M., K.S. (Kaori Sato), T.S., M.T., and K.O. performed experiments. K.S., H.S., W.K., K.N., T.S., T.Y., M.T., R.M., K.O., Y.O., T.C.S. and E.S. jointly analyzed and interpreted the data. K.S., H.S., W.K., K.N., T.S., T.Y., M.T., R.M., K.O., Y.O., T.C.S. and E.S. wrote and edited the manuscript. All authors provided feedback and agreed on the final manuscript.

## Acknowledgements

We thank Taisuke Tomita, University of Tokyo, for providing the antibodies to PS1. We thank Yukiko Nagai, RIKEN CBS, for secretarial assistance. Yuko Yamada, Tomoko Ishibuchi, Mitsuyoshi Togashi, Yoshihisa Sawada and members of Center of Basic Technology in Marmoset, Central Institute for Experimental Animals, are thanked for their technical assistance for producing and maintaining the AD models. We also thank Takayuki Mineshige and Terumi Yurimoto, Central Institute for Experimental Animals, for veterinary assistance. This work was supported by AMED under Grant Number JP20dm0207001 (Brain Mapping by Integrated Neurotechnologies for Disease Studies (Brain/MINDS)) (T.C.S.) and JP20dm0207065 (Study of developing neurodegenerative model marmosets and establishment of novel reproductive methodology) (E.S).

---

**Supplementary Figure 1.**
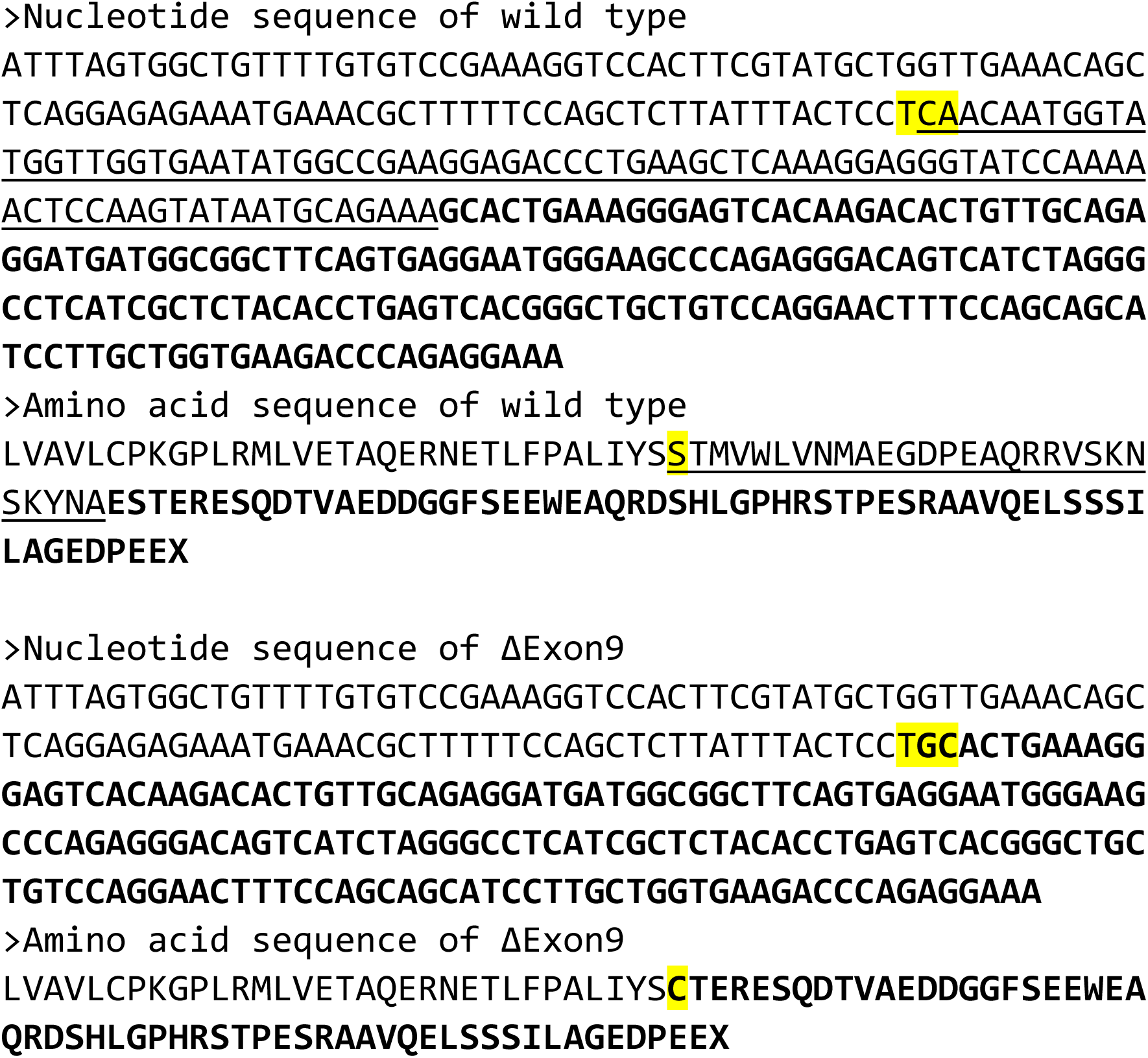
The cDNA and amino acid sequences of *PSEN1.* cDNA and amino acid sequences neighboring exon 9 of wild-type and mutant *PSEN1* mRNAs are shown. Normal characters: exon 8, underlined characters: Exon 9, bold characters: Exon 10, and characters highlighted in yellow: codons that encode for the 290th amino acid in the *PSEN1* gene. Note that TCA and TGC encode for serine (S) and cysteine (C), respectively.

**Supplementary Figure 2.**
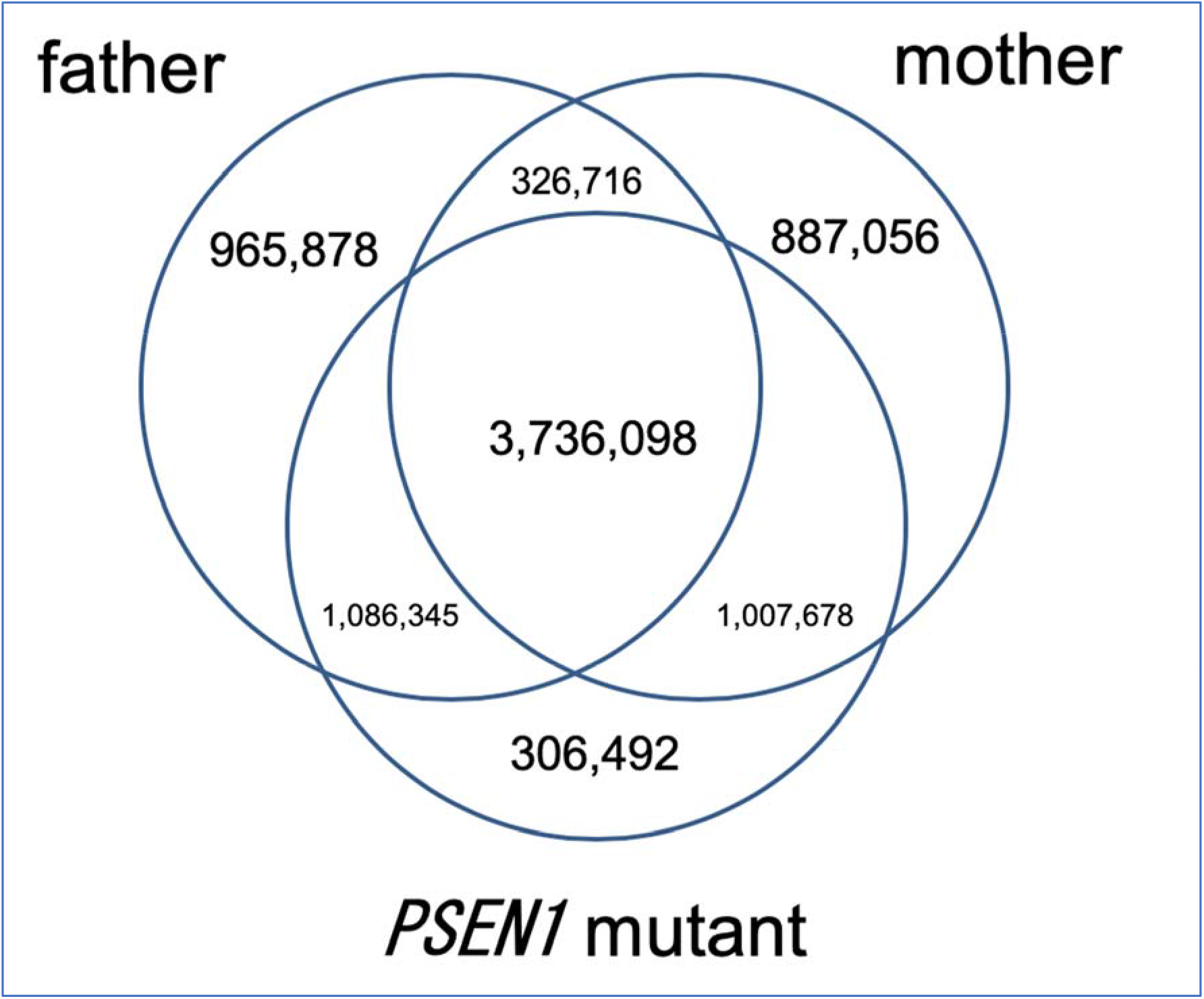
Variants identified in trio whole genome sequencing. Venn chart illustrating numbers and overlap of all variants identified in whole genome sequencing of the *PSEN1*-ΔE9 marmoset (I774gmF) and those from her parents. A highly permissive parameter in variant calling by Illumina DRAGEN pipeline (version 3.5.7) as a default setting was used, and no filtering was conducted. Of these all variants, 306,492 variants were identified in the *PSEN1*-ΔE9 marmoset (I774gmF) alone. None of the 306,492 variants were present within the 10 off-target candidate sites.

**Supplementary Table 1.**
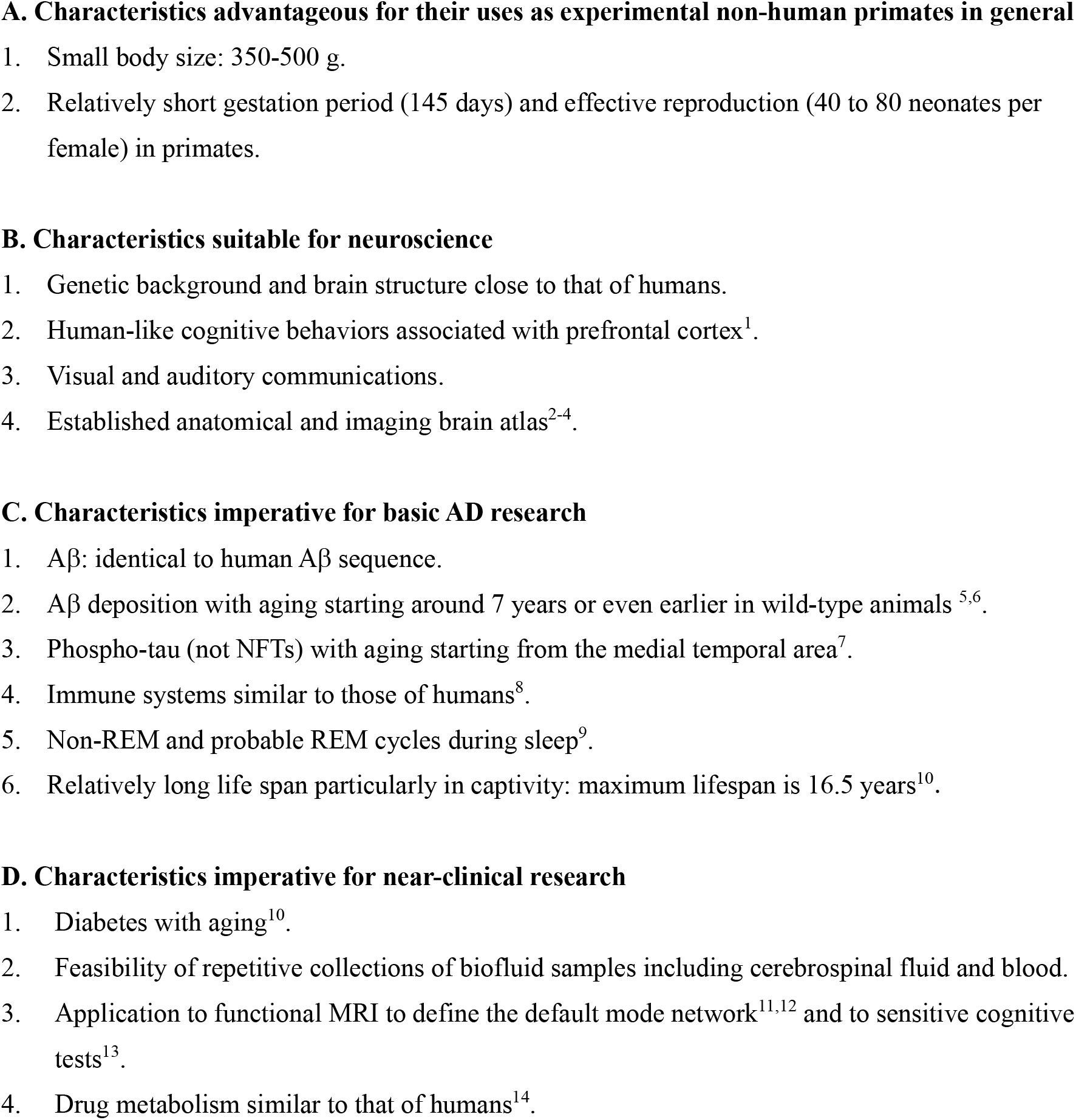
Merits of using marmosets for AD research.

**Supplementary Table 2.** Summary of off-target analysis.

| Off target | Chromosome | Position | Reference | Spacer length | TAL1 Score | TAL2 Score | Average Score | Mutation |
| --- | --- | --- | --- | --- | --- | --- | --- | --- |
| 1 | 21 | 7990751-7990801 | BLSI01000021.1 | 17 | 10.1 | 12.11 | 11.105 | None |
| 2 | 10 | 43948779-43948842 | BLSI01000010.1 | 30 | 15.79 | 6.66 | 11.225 | None |
| 3 | 6 | 70230528-70230582 | BLSI01000006.1 | 21 | 15.35 | 9.97 | 12.66 | None |
| 4 | 2 | 199375283-199375346 | BLSI01000002.1 | 30 | 12.75 | 13.18 | 12.965 | None |
| 5 | 2 | 33516787-33516844 | BLSI01000002.1 | 24 | 16.27 | 10.14 | 13.205 | None |
| 6 | 9 | 57942465-57942513 | BLSI01000009.1 | 15 | 11.81 | 14.72 | 13.265 | None |
| 7 | 13 | 91441037-91441093 | BLSI01000013.1 | 23 | 16.64 | 9.94 | 13.29 | None |
| 8 | 2 | 45917641-45917694 | BLSI01000002.1 | 20 | 12.52 | 14.3 | 13.41 | None |
| 9 | 13 | 61953631-61953684 | BLSI01000013.1 | 20 | 16.18 | 10.96 | 13.57 | None |
| 10 | 1 | 50319037-50319093 | BLSI01000001.1 | 23 | 16.1 | 11.16 | 13.63 | None |

**Supplementary Table 3.** Statistics of the whole genome sequencing data.

| Sample name | **father** | **mother** | ***PSEN1*-ΔE9** |
| --- | --- | --- | --- |
| Average sequenced coverage over genome (times over genome) | 57.41 | 62.19 | 42.07 |
| Total input reads [read] | 1,049,765,500 | 1,137,273,632 | 770,728,668 |
| Mapped reads | 1,043,044,330 | 1,130,059,433 | 765,829,950 |
| Unmapped reads | 6,721,170 | 7,214,199 | 4,898,718 |
| Paired reads (itself & mate mapped) | 1,039,914,838 | 1,126,580,296 | 763,452,884 |
| Reads with MAPQ [40:inf) | 982,059,798 | 1,064,082,451 | 714,555,792 |
| Reads with MAPQ [30:40) | 3,551,115 | 3,836,900 | 2,575,117 |
| Reads with MAPQ [20:30) | 5,595,307 | 6,015,031 | 4,192,941 |
| Reads with MAPQ [10:20) | 6,980,448 | 7,435,033 | 5,205,887 |
| Reads with MAPQ [ 0:10) | 44,857,662 | 48,690,018 | 39,300,213 |
| Reads with MAPQ NA (Unmapped reads) | 6,721,170 | 7,214,199 | 4,898,718 |
| Q30 bases [bp] | 145,168,078,063 | 156,463,755,481 | 106,214,269,234 |

**Supplementary Table 4.**
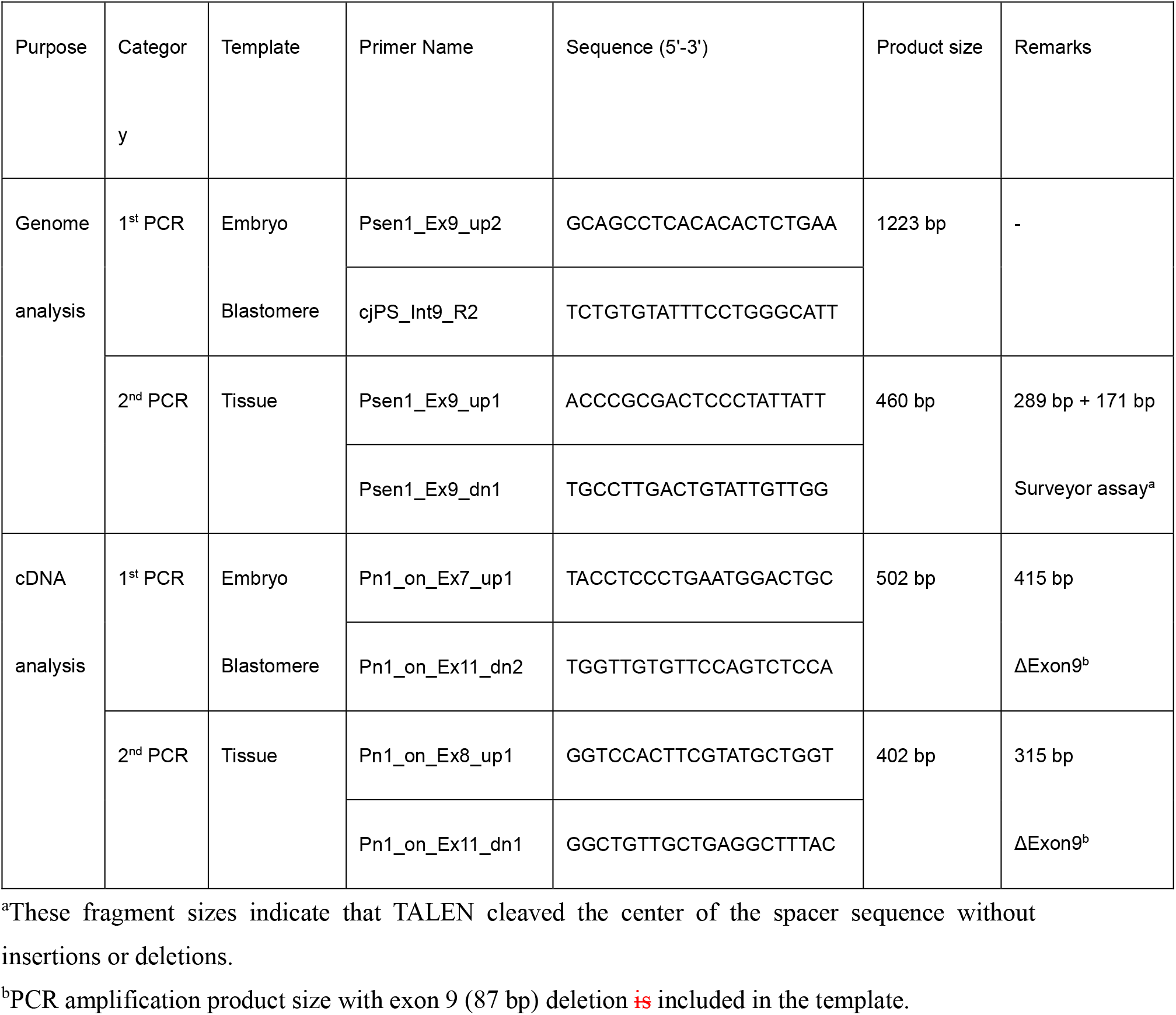
Primers used in the present study.

**Supplementary Table 5.**
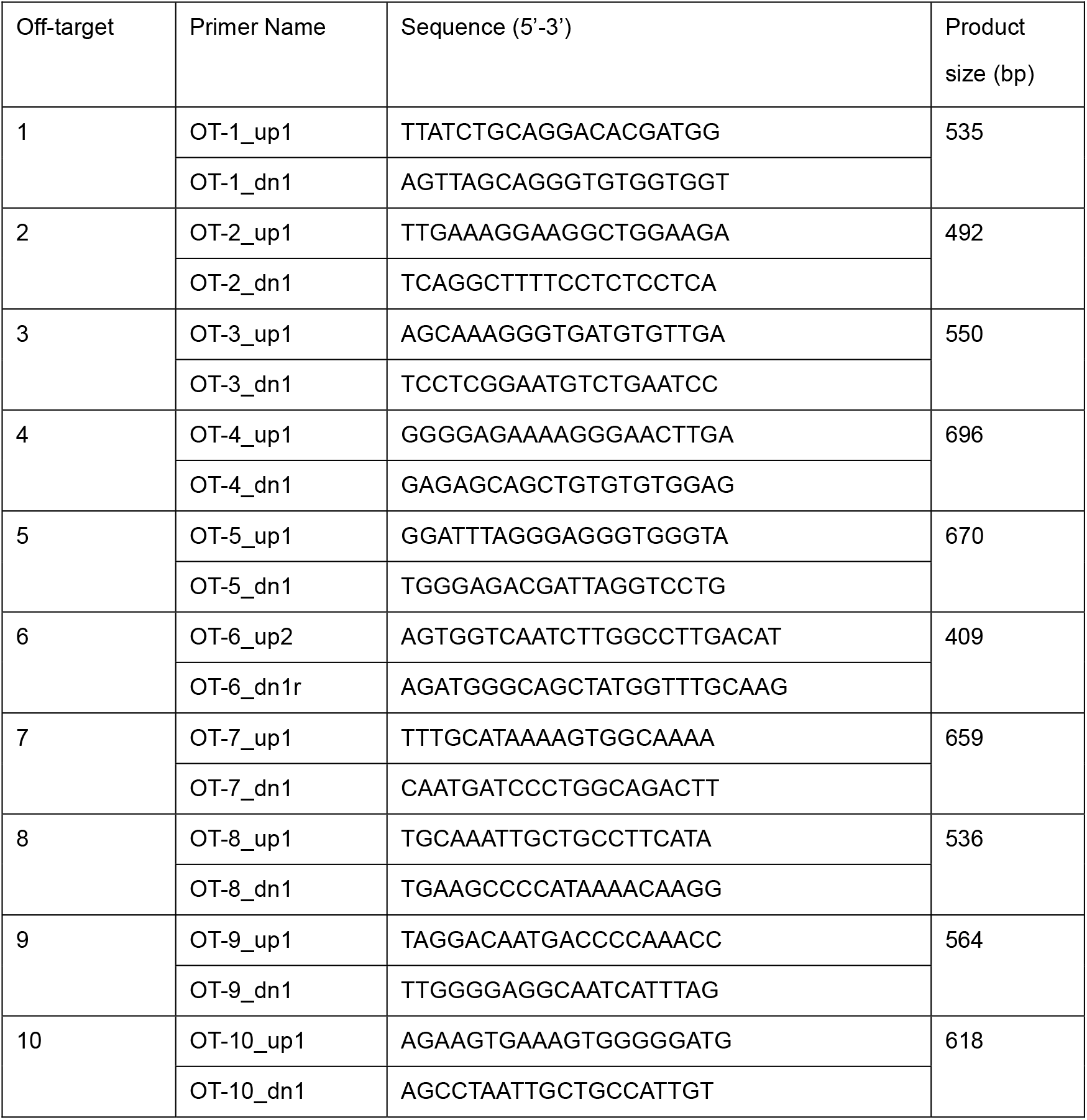
Primers used for the off-target analysis.

